# Ultrastructure analysis of mitochondria, lipid droplet and sarcoplasmic reticulum apposition in human heart failure

**DOI:** 10.1101/2025.01.29.635600

**Authors:** Nadina R. Latchman, Tyler L. Stevens, Kenneth C. Bedi, Benjamin L. Prosser, Kenneth B. Margulies, John W. Elrod

## Abstract

**Background:** Cardiomyocyte structural remodeling is reported as a causal contributor to heart failure (HF) development and progression. Growing evidence highlights the role of organelle apposition in cardiomyocyte function and homeostasis. Disruptions in organelle crosstalk, such as that between the sarcoplasmic reticulum (SR) and mitochondria, are thought to impact numerous cellular processes such as calcium handling and cellular bioenergetics; two processes that are disrupted and implicated in cardiac pathophysiology. While the physical distance between organelles is thought to be essential for homeostatic cardiomyocyte function, whether the interactions and coupling of organelles are altered in human heart failure remains unclear.

**Methods:** Here, we utilized transmission electron microscopy and careful quantification of ultrastructure to characterize the changes in organelle apposition in cardiomyocytes isolated from the hearts of patients diagnosed with various types of HF. Subsequently we employed molecular approaches to examine the expression of proposed organelle tethers.

**Results:** We demonstrate that cardiomyocytes isolated from dilated cardiomyopathy, hypertrophic cardiomyopathy and ischemic cardiomyopathy hearts display smaller, more rounded mitochondria, as compared to nonfailing controls. Failing cardiomyocytes also exhibited disrupted SR-mitochondria juxtaposition and changes in the expression of proposed molecular tethers. Further analysis revealed alterations in lipid droplet dynamics including decreased lipid droplet content and less lipid droplets in association with mitochondria in failing cardiomyocytes.

**Conclusion:** Here we observed changes in organelle dynamics in cardiomyocytes isolated from heart failure patients diagnosed with differing etiologies. Our results suggest that organelle structure and apposition may be a ubiquitous contributor to human HF progression.

**RESEARCH PERSPECTIVE:** **What is New?**

- We provide a detailed analysis of organelle apposition in human heart failure, which has been understudied, and report that that failing human cardiomyocytes display an increase in distance between mitochondria and both the sarcoplasmic reticulum and lipid droplets.
- Structural changes in organelles are correlated with the expression of proposed organelle tethers.
- Resource of ultrastructural changes in organelle apposition in human heart failure resulting from various etiologies.

**What Questions Should be Addressed next?**

- The results from this study provide rationale for causal experimentation to elucidate the contribution of organelle apposition to the progression of heart failure. Future studies examining mechanisms of mitochondrial tethering to the SR or lipid droplet will evaluate specific targets for therapeutic application.

## INTRODUCTION

Heart failure (HF) is a leading cause of morbidity and mortality worldwide with an increasing prevalence^1–3^. Despite its complex etiology, a common feature of HF is the heart’s inability to pump blood to meet the body’s metabolic needs^3^. This process relies on the contraction of cardiomyocytes, highly energy-dependent cells within the heart, to ensure appropriate cardiac function^4–6^. Cardiac contractions are driven by the highly specialized and intricate organization of cardiomyocytes, including dynamic organelle interactions within the cell^6–8^. Alterations in cardiomyocyte structure and function, such as transverse tubule (T-tubule) remodeling, sarcomere disorder, and impaired mitochondrial dynamics can disrupt cardiac homeostasis and are implicated in heart failure pathogenesis resulting from diverse etiologies^6–16^.

In cardiomyocytes, organelles cooperate to mediate the cellular processes necessary for cardiac homeostasis with accumulating evidence suggesting that organelle communication is critical for maintaining cardiac function^5–7,15–23^. The significance of organelle interplay and juxtaposition is highlighted by the proximity of T-tubules and the junctional sarcoplasmic reticulum (jSR) which is necessary for excitation-contraction coupling (ECC)^4–8^. Mitochondria situated near the SR take up calcium to couple energetic demand with the massive workload of the heart^15–23^. SR-mitochondria crosstalk has also been shown to play a pivotal role in bioenergetics, calcium homeostasis, oxidative stress, mitochondrial dynamics, cell survival, lipid metabolism, and lipid homeostasis^16,22–38^. Emerging evidence has also highlighted the critical role of interactions between lipid droplets (LD) and mitochondria within cardiomyocytes to maintain energy homeostasis, lipid composition, and metabolic regulation^33–38^. Therefore, interaction and communication between organelles enable cardiomyocytes to adapt to their dynamic environments. Inducing heart failure in various murine models revealed ultrastructural remodeling of SR-mitochondria contacts^39–43^. Furthermore, the crosstalk (physical distance) between the SR and mitochondria is disrupted. This leads to progressive declines in cardiomyocyte function, impacting several critical functions such as the maintenance of calcium (Ca^2+^) homeostasis, bioenergetics, lipid metabolism, protein homeostasis, cell fate, cell death, and autophagy^39–48^.

The physical proximity and functional interplay between organelles are established by molecular tethers^49–58^. These specialized protein tethers bring organelles in close apposition to each other without fusion of individual membranes, enabling organelle communication, trafficking, and exchanges of lipids and ions^51–57^. Seminal electron microscopy ultrastructure studies revealed that protein tethers scaffold organelles at approximately 10-30 nm^53–57^. Membrane tethering can be achieved by single proteins or oligomeric protein complexes, with the list of proteins proposed to mediate these contact sites continuing to increase^51–62^. Notable tethers proposed to mediate SR-mitochondria contacts include mitofusin 2 (MFN2), vesicle-associated membrane protein-associated protein B (VAPB), glucose regulated protein 75 (GRP75), protein tyrosine phosphatase-interacting protein 51 (PTPIP51) and FUN14 domain-containing protein 1 (FUNDC1), while proteins proposed to mediate LD-mitochondria interactions include Perilipin-5 (PLIN5) and Mitoguardin-2 (MIGA2)^63–77^. However, the detailed roles of these proposed membrane tethers remain controversial as most all have a diverse range of functions and are not merely ‘professional’ tethers.

Although understanding the pathophysiological role of organelle apposition in cardiovascular disease has progressed in recent years, the exact mechanisms of organelle apposition, specifically how SR-mitochondria and LD-mitochondria tethering contributes to human heart failure, remains poorly characterized. To define the changes in organelle apposition in human heart failure, we utilized transmission electron microscopy (TEM) to evaluate mitochondrial and lipid droplet ultrastructure, as well as to investigate SR-mitochondria and LD-mitochondria association. In addition, we examined the expression of several proposed organelle tethers to further correlate these findings. Here, we report altered mitochondria and lipid ultrastructure in failing human cardiomyocytes. Failing cardiomyocytes exhibited decreased size of both mitochondria and lipid droplets. Failing cardiomyocytes also had rounder mitochondria compared to nonfailing controls. In addition, we observed increased SR-mitochondria distance, which correlated with alterations in mRNA tether expression. Decreased LD-mitochondria association in failing cardiomyocytes was also observed. Overall, this data suggests that alterations in organelle apposition may contribute to HF progression.

## METHODS

### Human Left Ventricular Samples

Human myocardium was obtained from end-stage failing hearts explanted at time of cardiac transplantation. Non-failing donor hearts that were deemed unsuitable for transplantation were used as control. Patient consent, sample collection and preparation, and clinical data collection were all performed according to a protocol approved by the Lewis Katz School of Medicine Institutional Review Board

### Transmission electron microscopy sample preparation

Left ventricular samples were dissected into 2% glutaraldehyde and 2% paraformaldehyde in 0.1M sodium cacodylate buffer and cut into small pieces, approximately 1mm3, and held in fixative at 4°C until processing could be completed. Samples were washed in 0.1M sodium cacodylate buffer pH 7.4, three times 15 min each wash, and postfixed in freshly prepared 1% osmium tetroxide and 1.5% potassium ferrocyanide in 0.1M sodium cacodylate buffer pH 7.4 for two hours. The tissues were washed in Nanopure water, four times 15 min each, and then stained in 1% uranyl acetate overnight at room temperature, protected from light. Following three washes with Nanopure water, 15 min each wash, the tissues were dehydrated using an ascending series of acetone solutions (25% acetone, 50% acetone, 75% acetone, 95% acetone, 100% glass-distilled acetone, 100% glass-distilled acetone), 15 min each step, and then were infiltrated with increasing concentrations of Spurr low viscosity resin (25% Spurr resin in acetone, 50% Spurr resin in acetone, 75% Spurr resin in acetone), one hour each step on a rotator. Following two exchanges with 100% Spurr resin, one hour each, samples were infiltrated with 100% Spurr resin overnight on a rotator at room temperature. The next day, tissues were embedded in aluminum foil dishes with freshly prepared Spurr resin and polymerized at 60°C for 48h. Tissue pieces exhibiting proper orientation were excised from the polymerized resin using a jeweler’s saw and glued onto a BEEM capsule support using cyanoacrylate glue. Blocks were sectioned on a Leica UC7 ultramicrotome, and ultrathin sections were collected onto 200 mesh formvar/carbon coated grids. Grids were stained with Reynolds lead citrate and 1% uranyl acetate in 50% methanol. Grids were examined and imaged in a FEI Tecnai 12 120 keV digital TEM, with images acquired at various magnifications (e.g. 1,100x – 6500x).

### Transmission electron microscopy morphometric analysis

TEM morphometric analysis of mitochondria, lipid, and mitochondria-SR associations was performed using ImageJ/FIJI (NIH). After calibration for distance, mitochondrial shape descriptors and size measurements were obtained utilizing ImageJ (NIH) by manually tracing only discernable mitochondria. Circularity is computed as [4πx surface area)], roundness [4*(surface area)/(πx major axis^2^) these two indexes of sphericity with values of 1 indicate perfect spheres and Feret Diameter represents the longest distance between any two points within a given mitochondrion^78^. For mitochondria-SR associations and LD-Mitochondria association MitoCare Plugin for image J was utilized^41^. Areas where SR was <50nm from OMM were determined as SR-Mitochondrion interface. To obtain the mean gap distance, mitochondrion was first traced followed by tracing SR. Areas where the LD was <100nm from the outer mitochondrial membrane (OMM) were determined as a LD-mitochondrion interface. To obtain mean gap distance, the LD membrane was first traced followed by tracing the mitochondrion OMM with values obtained from the plugin Raw data were analyzed by Prism 9 (Graph Pad Software)^41,78^.

### qPCR gene expression analysis

RNA was isolated using Qiagen RNeasy Fibrous Tissue Kit (Qiagen, 74704) according to the manufacturer’s instructions. cDNA was generated using the Applied Biosystems High-Capacity cDNA Reverse Transcription Kit (Applied Biosystems, 4368814). Quantitative PCR (qPCR) was performed on a CFX96 Touch Real-Time PCR Detection System (Bio-Rad) using PowerUp SYBR Green Master Mix (Applied Biosciences #100029283). The real-time PCR conditions were: UDG activation at 50°C for 2 minutes, initial denaturation at 95°C for 10 minutes, followed by 40 cycles (95°C for 15 seconds, 60°C for 30 seconds, 72°C for 30 seconds). qPCR primers against human transcripts are listed in Supplemental Table 1.

### Statistics

All results are presented as mean ± SEM. Statistical analyses were used Prism 9.0 (Graph Pad Software) and Microsoft Excel. Statistical parameters including sample size and statistical significance values are included in each figure legend. All statistical tests compared NF to each HF etiology. Column analyses were performed using a one-way ANOVA with Dunnett’s post-hoc analysis. For grouped analyses, a two-way analysis of variance (ANOVA) with Dunnett’s post-hoc analysis was performed. For categorical data analyzing counts in different distance bins, Chi-square analysis was performed. For all analyses, *P* values less than 0.05 were considered significant.

## RESULTS

### Failing Cardiomyocytes Exhibit Altered Mitochondrial Dynamics

Mitochondrial morphology is essential in maintaining the physiological function of cardiomyocytes. Alterations in mitochondrial structure induce mitochondria dysfunction, resulting in reduced energy production and decreased contractility in cardiomyocytes^12–18,78–82^. A previous study has reported that decreased mitochondrial size is correlated with impaired bioenergetics^83^. To characterize mitochondria ultrastructure in human heart failure (HF), we obtained left ventricular tissue from nonfailing hearts (NF), as well as hearts with dilated cardiomyopathy (DCM), hypertrophic cardiomyopathy (HCM) or ischemic cardiomyopathy (ICM). The characteristics of the patients at the time of sample collection are depicted in Table 1, with patient demographic information being provided in Table 2. Characterization of heart failure etiology was determined by clinical features. Nonfailing controls were from healthy hearts not suitable for translation with a left ventricular ejection fraction ≥ 50%. Dilated cardiomyopathy were from hearts that required transplantation not due to ischemic cardiomyopathy. Ischemic cardiomyopathy were hearts from patients that required transplantation due to severe coronary artery disease. Hypertrophic cardiomyopathy was determined by both echocardiography and magnetic resonance imaging. We performed TEM on LV samples to evaluate mitochondrial size and morphology (Figure 1A-B). While there were no significant differences in overall mitochondrial area, DCM and HCM samples displayed an increased number of mitochondria per cardiomyocyte area, indicating increased fragmentation and disruption in the balance of fission and fusion dynamics (Figure 1C-D, S1A-B). This was confirmed by failing cardiomyocytes exhibiting ultrastructure alternations, including significant reductions in Feret diameter and mitochondrial perimeter (Figure 1E-F, S1C-D). An increased frequency of smaller mitochondria was identified in failing vs. nonfailing cardiomyocytes (Figure 1G, Table 3), further supporting increased mitochondrial fragmentation in heart failure. Topographical analysis revealed that cardiomyocytes of failing hearts demonstrated a significant decrease in the average mitochondria perimeter per patient (Figure S1E). Patients with DCM had ∼13% decrease in mitochondria perimeter compared to NF controls while ICM and HCM hearts demonstrated a ∼10% decrease (Figure S1F). Analysis of mitochondria morphology revealed that cardiomyocytes from each HF etiology had significantly increased mitochondrial circularity compared to nonfailing controls, suggesting a damaged and fragmented mitochondrial population (Figure 1H, S1G). Changes in mitochondrial morphology, quantity, and position are referred to as mitochondrial dynamics, with disrupted dynamics being associated with perturbed mitochondrial function cardiomyocytes^12–18,78–84^. To determine if transcriptional changes of any regulators of mitochondrial dynamics correlate with altered mitochondrial morphology from our TEM analysis, we performed mRNA expression analysis on LV patient samples (Figure 1I-M). While there were no consistent alterations in the expression of mitochondrial dynamic regulators in each type of HF, we did observe a significant upregulation of *OPA1* in ICM hearts (Figure 1J), as well as an upregulation of *ERMIN2,* the splice variant of *MFN2* reported to regulate ER morphology^85^, in HCM hearts (Figure 1M). Together, this ultrastructure analysis reveals that failing cardiomyocytes have smaller and more fragmented mitochondria, indicative of severe changes in mitochondrial dynamics and likely dysfunction.

**Figure 1.**
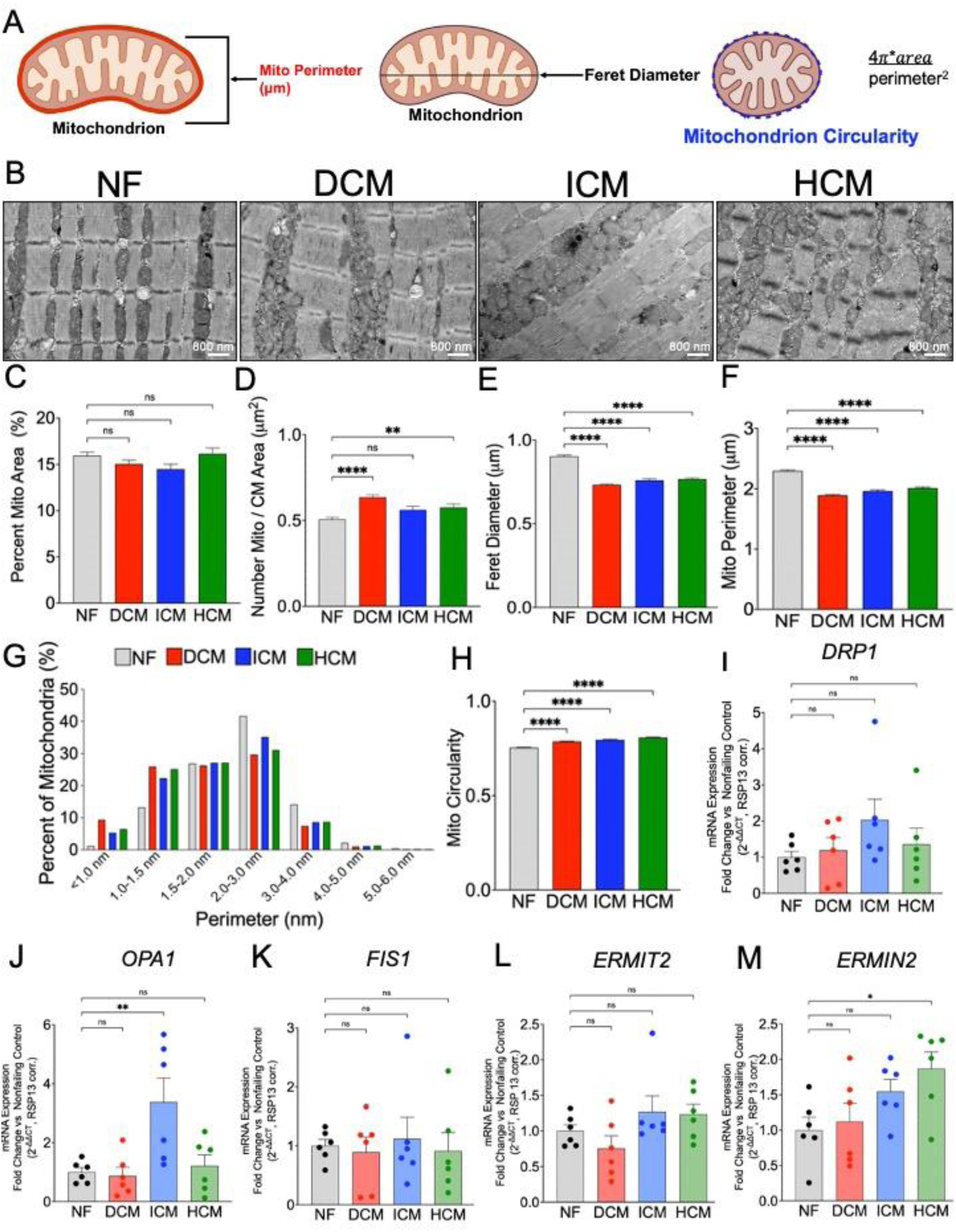
Mitochondrial dynamics are altered in failing human cardiomyocytes. (A) Schematic for mitochondrial ultrastructure analysis. (B) Representative EM Images. Cardiac tissue from NF, DCM, HCM, ICM samples were sectioned and imaged by TEM. TEM images were analyzed and quantified using ImageJ Fiji (Scale bars: 800nm). Ultrastructure measurements include (C) percent mitochondria area, (D) number of mitochondria/cardiomyocyte (CM) area, (E) mitochondria Feret diameter, (F) mitochondria perimeter, (G) frequency of mitochondrial perimeter in HF etiologies, and (H) mitochondrial circularity. qPCR analysis of genes involved in mitochondrial dynamics including *DRP1*, (J) *OPA1*, (K) *FIS1*, (L) *ERMIT2*, (M) and *ERMIN2*. Number of patients for EM Data=8 NF, 9 DCM, 3 ICM, 5 HCM. Number of mito= 2760 NF, 4546 DCM, 1368 ICM, 2350 HCM. Number of patients for qPCR = 6 NF, 6 DCM, 6 ICM, 6HCM. Individual data points presented in supplemental. Data is presented as mean ± SEM. ****p<0.0001, **p<0.01, *p<0.05; One-way ANOVA with Dunnett’s Post Hoc Test. Abbreviations: EM: Electron Microscopy, HF- Heart Failure, NF-Nonfailing, DCM- Dilated Cardiomyopathy, ICM- Ischemic Cardiomyopathy, HCM- Hypertrophic Cardiomyopathy, TEM-Transmission Electron Microscopy, *DRP1- Dynamin-related protein* 1*OPA1- OPA1 Mitochondrial Dynamin Like GTPase, FIS1- Fission, Mitochondrial 1, ERMIT2- ER mitofusin 2 tether, ERMIN2-ER Mitofusin 2*.

**Table 1:**
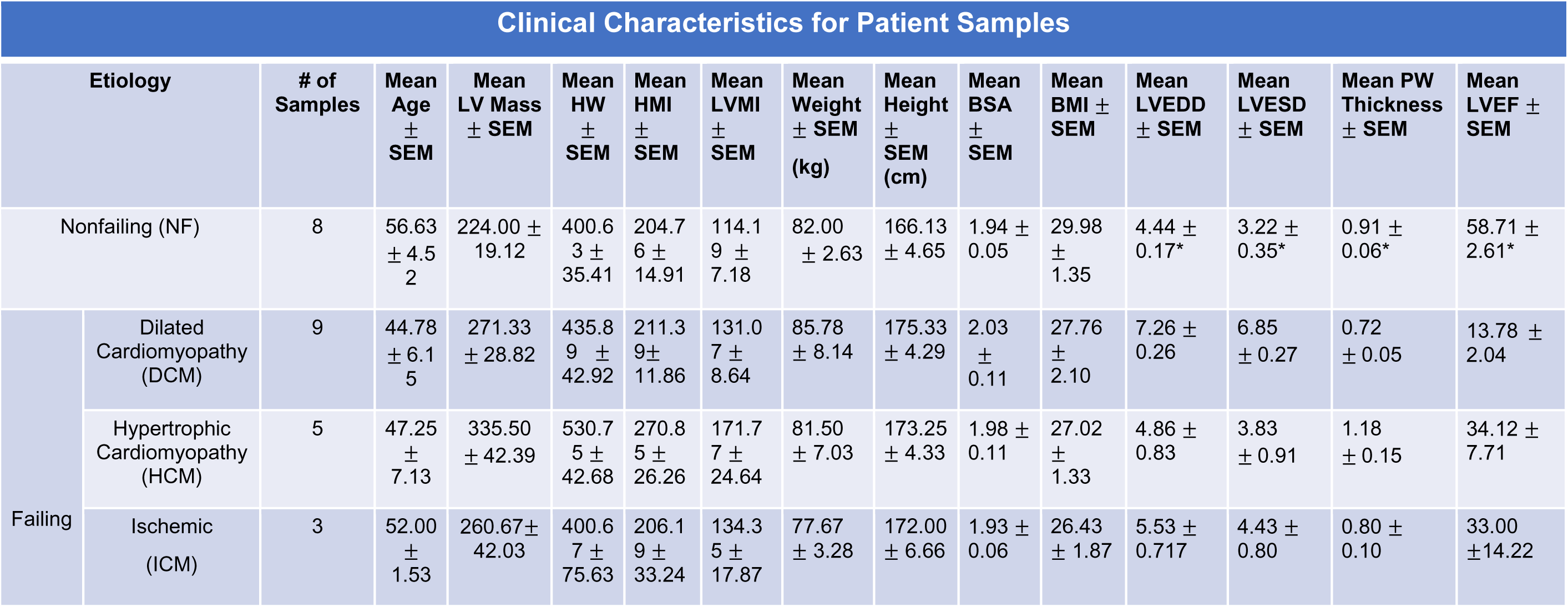
Clinical characteristics of patients from different heart failure etiologies used in Figures 1-4. Abbreviations: LV- Left ventricular, HW- Heart Weight, HMI-Heart Mass Index, LVMI-Left Ventricular Mass Index, kg-kilogram, cm- centimeters, BMI-Body Mass Index, LVEDD-Left Ventricular End Diastolic Diameter, LVESD- Left Ventricular End Systolic Diameter, PW- Posterior Wall, LVEF- Left Ventricular ejection fraction

**Table 2:**
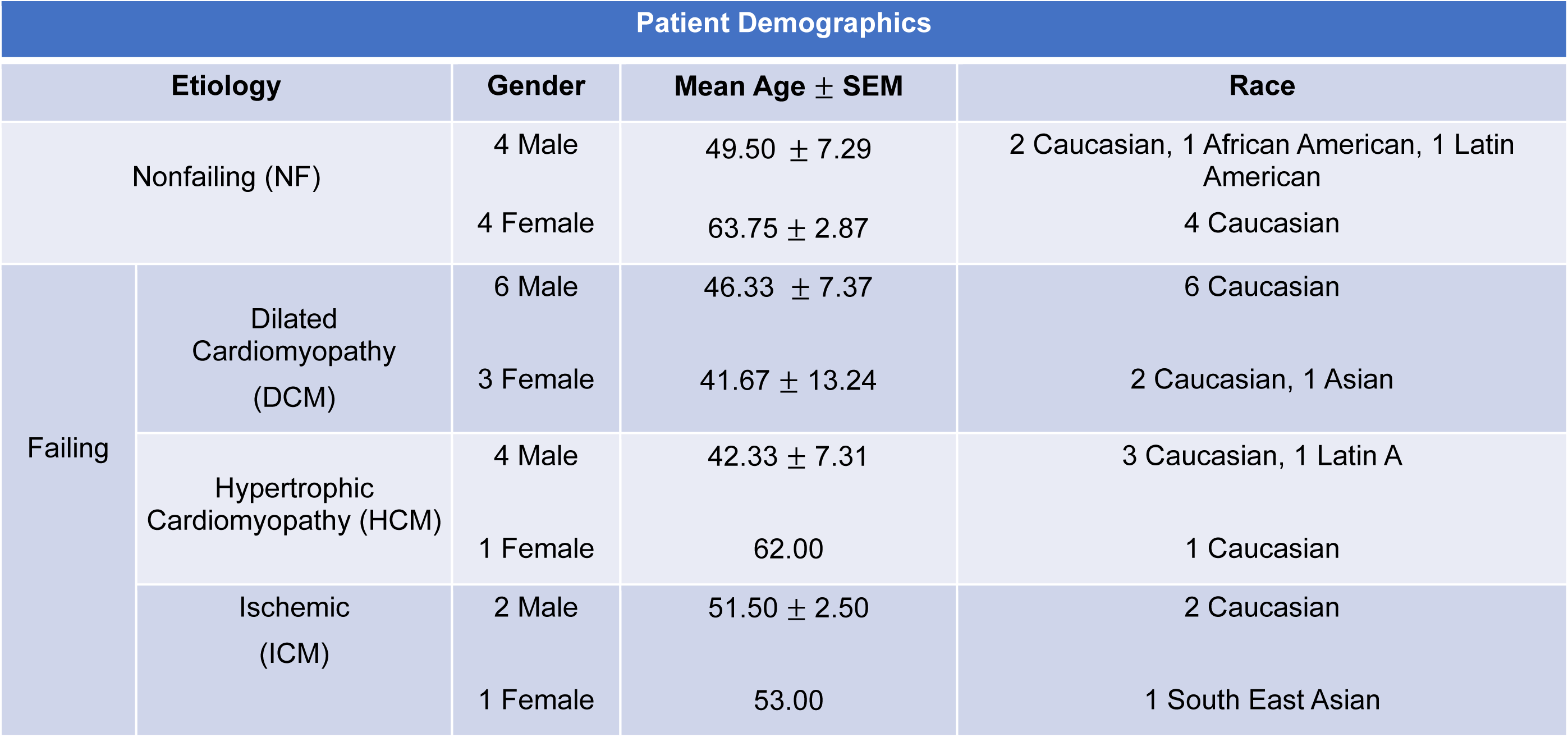
Patient demographics from different heart failure etiologies used in Figures 1-4.

**Table 3:**
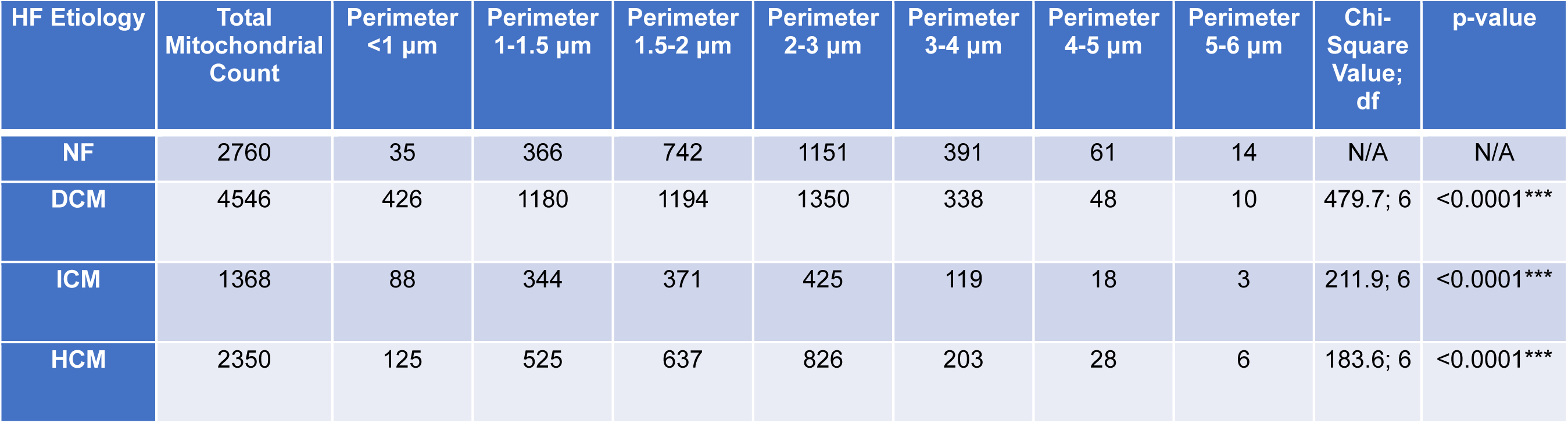
Mitochondrial count and mitochondrial perimeter size from different heart failure etiologies used in Figure 1. Chi-Square value and df comparing non-failing group to corresponding HF etiology. Abbreviations: HF- Heart Failure, NF- Nonfailing, DCM- Dilated Cardiomyopathy, ICM- Ischemic Cardiomyopathy, HCM- Hypertrophic Cardiomyopathy.

### SR-Mitochondria Distance is Increased in Failing Cardiomyocytes

In cardiomyocytes, SR and mitochondria form close contacts to carry out ECC, calcium signaling, and lipid transfer^15–38^. Disruptions in SR-mitochondria contacts have been implicated in heart failure progression^39–48^. We utilized TEM analysis to examine healthy and heart failure LV samples to determine if alterations in mitochondria ultrastructure had any impact on SR-mitochondria apposition (Figure 2A-B). Analysis revealed that cardiomyocytes from DCM, ICM and HCM patient hearts displayed a significant increase in mean and minimum SR-mitochondria distances compared to nonfailing controls, indicating an increase in the overall distance between the SR and mitochondria (Figure 2C-2D, S2A-B). Additionally, the length of outer mitochondrial membrane in contact with the SR is a strong indicator of functional interaction between the SR and mitochondria. Here, we observed that failing cardiomyocytes had both a decrease in the length of the SR within a 50-nm distance from the outer mitochondrial membrane (OMM), as well as a decreased length of OMM in association with SR at distance of ≤ 50nm (Figure 2E-F, S2C-D). Interestingly, nonfailing cardiomyocytes had significantly more SR-mitochondria contacts within <20 nm proximity (Figure 2G-H, S2E, Table 4). Minimal changes to the number of SR-Mito contacts were observed between DCM and nonfailing cardiomyocytes. However, there was a significant decrease in the number of SR-mitochondria contacts in HCM cardiomyocytes, along with a trending decrease in ICM cardiomyocytes (Figure 2I, S2F). Taken together, these data suggest that there is substantially diminished physical interaction between the SR and mitochondria in failing cardiomyocytes compared to nonfailing controls.

**Figure 2.**
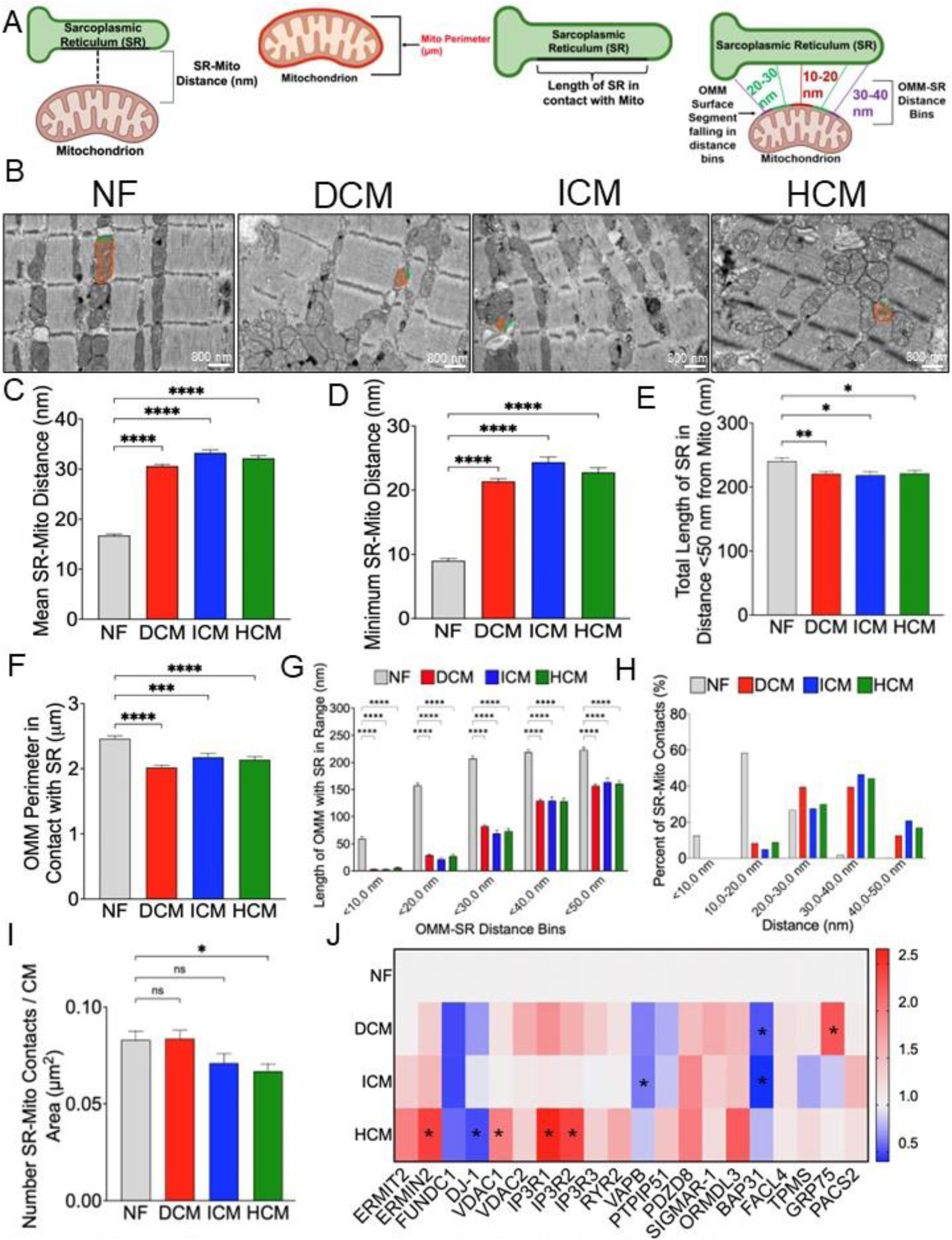
Failing cardiomyocytes have altered sarcoplasmic reticulum-mitochondria apposition. (A) Schematic of mitochondria-sarcoplasmic reticulum apposition analysis. (B) Representative EM Images. Cardiac tissue from NF, DCM, HCM, ICM samples were sectioned and imaged by TEM. TEM images were analyzed and quantified using ImageJ Fiji (Scale bars: 800nm). Examples of mitochondria-SR interactions are highlighted in orange and green respectively. Ultrastructure measurements include (C) mean SR-mitochondria distance, (D) minimum SR-mitochondria distance, (E) total length of SR less than 50 nm from mitochondria, (F) mitochondrial perimeter in association with SR, (G) binning of the length of OMM less than 50 nm from SR, (H) frequency of SR-mito contacts, (I) and total number of SR-mitochondrial contracts per image area. (J) Heatmap representation of mRNA levels of tethers in HF depicted as fold change corrected to calnexin. Number of patients=8 NF, 9 DCM, 3 ICM, 5 HCM Number of contacts=405 NF, 526 DCM, 163 ICM, 237 HCM. Number of patients for qPCR = 6 NF, 6 DCM, 6 ICM, 6HCM. Individual data points presented in supplemental. Data is presented as mean ± SEM. ****p<0.0001, **p<0.01, *p<0.05; One-way ANOVA with Dunnett’s Post Hoc Test unless stated otherwise. 2G Two-Way ANOVA with Dunnett’s Post Hoc Test. Abbreviations: EM: Electron Microscopy, HF- Heart Failure, NF-Nonfailing, DCM- Dilated Cardiomyopathy, ICM- Ischemic Cardiomyopathy, HCM- Hypertrophic Cardiomyopathy, TEM-Transmission Electron Microscopy, SR- Sarcoplasmic Reticulum, OMM- Outer Mitochondria Membrane.

**Table 4:**
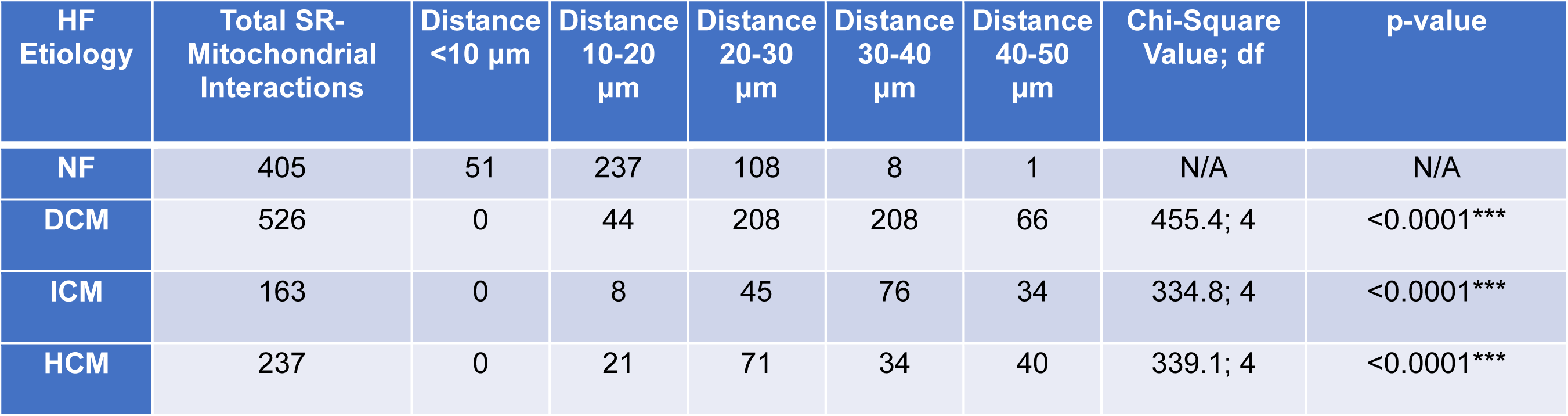
SR-Mitochondrial interaction counts binned by average distance from different heart failure etiologies used in Figure 2. Chi-Square value and df comparing non-failing group to corresponding HF etiology. Abbreviations: HF- Heart Failure, NF-Nonfailing, DCM- Dilated Cardiomyopathy, ICM- Ischemic Cardiomyopathy, HCM- Hypertrophic Cardiomyopathy, SR- Sarcoplasmic Reticulum.

Several proteins have been implicated in the regulation of the SR-mitochondria tethering complex ^64–69,85^. To corroborate the structural and topographical findings observed in TEM analysis, qPCR analysis was performed on tissue from human left ventricular tissue to evaluate the expression of proposed SR-mitochondria tethers (Figure 2J). We found decreased mRNA expression of *VAPB* and *BAP31* in both DCM and ICM compared to nonfailing LV samples, in addition to increased *GRP75* expression exclusively in DCM. Interestingly, RNA isolated from HCM hearts exhibited the most notable changes in tethering components, with decreased expression of *FUNDC1* and *DJ1* and increased expression of the Mitofusin 2 splice variant, *ERMIN2*, as well as *VDAC1* and *IP3R* (Figure 2J). Overall, these results suggest that the observed disruption in SR-mito juxtaposition may result from transcriptional remodeling, and this may contribute to the impaired cardiomyocyte homeostasis contributing to HF progression.

### Lipid Droplet dynamics are altered in failing cardiomyocytes

Lipid droplets (LDs) are dynamic organelles that play a role in lipid metabolism and energy homeostasis and LD structure/function is proposed to be disrupted in heart failure^34–36,86^. To gain a deeper understanding of the role of LDs in heart failure development, we performed LD ultrastructure analysis (Figure 3A-B). TEM analysis revealed that failing cardiomyocytes had a significant reduction in both the total area and number of LDs per cardiomyocyte area (Figure 3C-D, S3A-B), suggesting overall diminished lipid storage and/or increased utilization. Failing cardiomyocytes displayed a decrease in LD perimeter compared to NF cardiomyocytes (Figure 3E, S3C). We binned LD samples by perimeter size, and cardiomyocytes from DCM LV samples had a significantly downward shift in LD perimeter in addition to a trending downward shift ICM cardiomyocytes (Figure 3F, Table 5). A significant decrease in Feret diameter was also observed in failing cardiomyocytes across all etiologies (Figure 3G, S3D). Despite changes in LD size, TEM analysis did not reveal any significant changes in LD circularity (Figure 3H, S3E). Overall, ultrastructure analysis identified failing cardiomyocytes have fewer LDs that are smaller in size, which may have several implications, including increased lipolysis, impaired lipid storage, and/or dysregulation of lipid metabolism.

**Figure 3.**
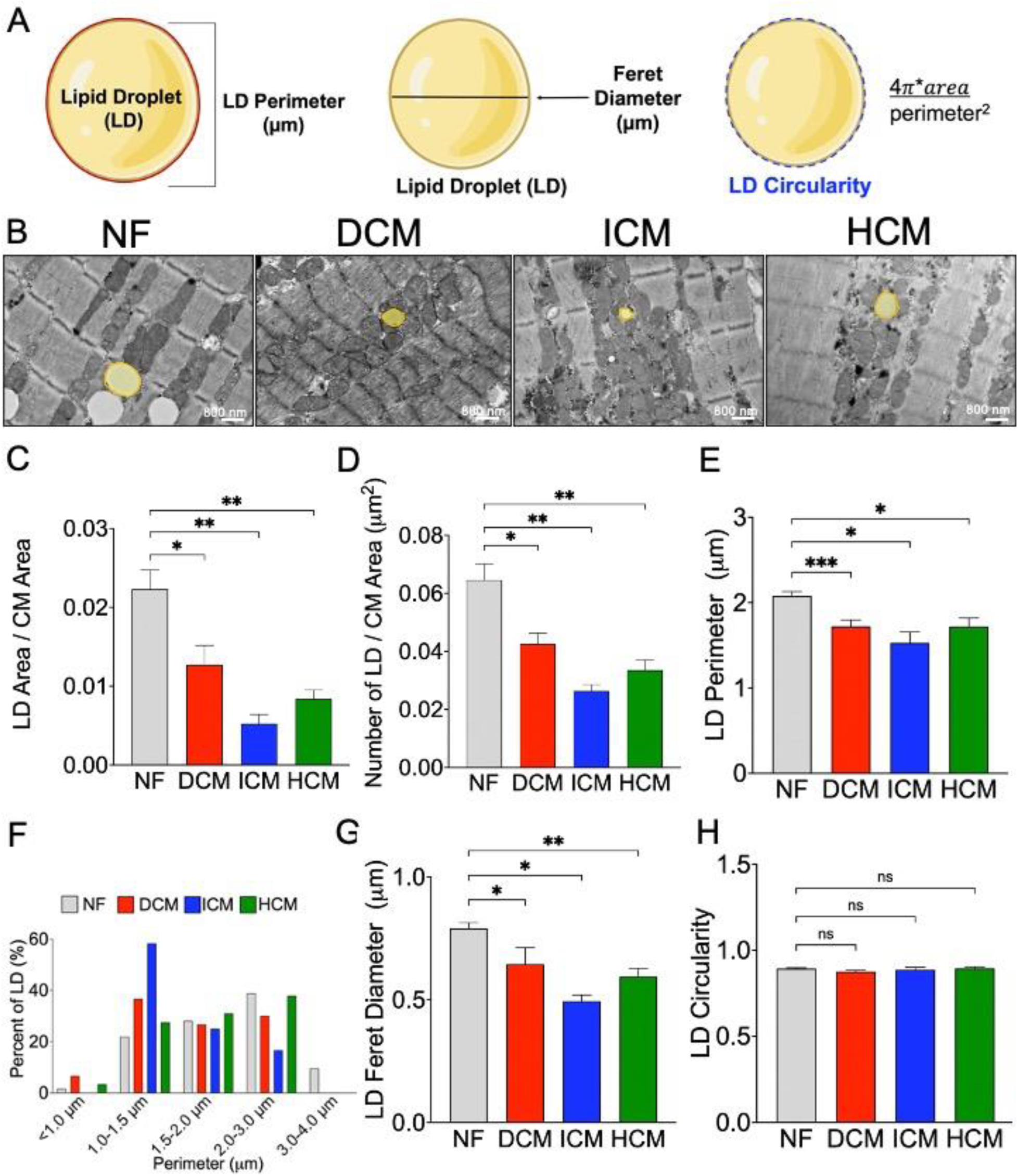
LD ultrastructure is altered in failing cardiomyocytes. (A) Schematic of lipid droplet (LD) ultrastructure analysis. (B) Representative EM Images. Cardiac tissue from NF, DCM, HCM, ICM samples were sectioned and imaged by TEM. TEM images were analyzed and quantified using ImageJ Fiji (Scale bars: 800nm). Ultrastructure measurements include, (C) LD area/total cardiomyocyte area, (D) number of LD/CM area, (E) LD perimeter, (F) Percentage of LD perimeter sizes per HF etiology, (G) LD Feret diameter, (H) and LD Circularity. Number of patients=8 NF, 9 DCM, 3 ICM, 5 HCM Number of LD= 178 NF, 60 DCM, 12 ICM, 29 HCM. Individual data points presented in supplemental. Data is presented as mean ± SEM. **p<0.01, *p<0.05; One-way ANOVA with Dunnett’s Post Hoc Test. Abbreviations: EM: Electron Microscopy, HF- Heart Failure, NF-Nonfailing, DCM- Dilated Cardiomyopathy, ICM- Ischemic Cardiomyopathy, HCM- Hypertrophic Cardiomyopathy, TEM-Transmission Electron Microscopy, CM- Cardiomyocyte.

**Table 5:**
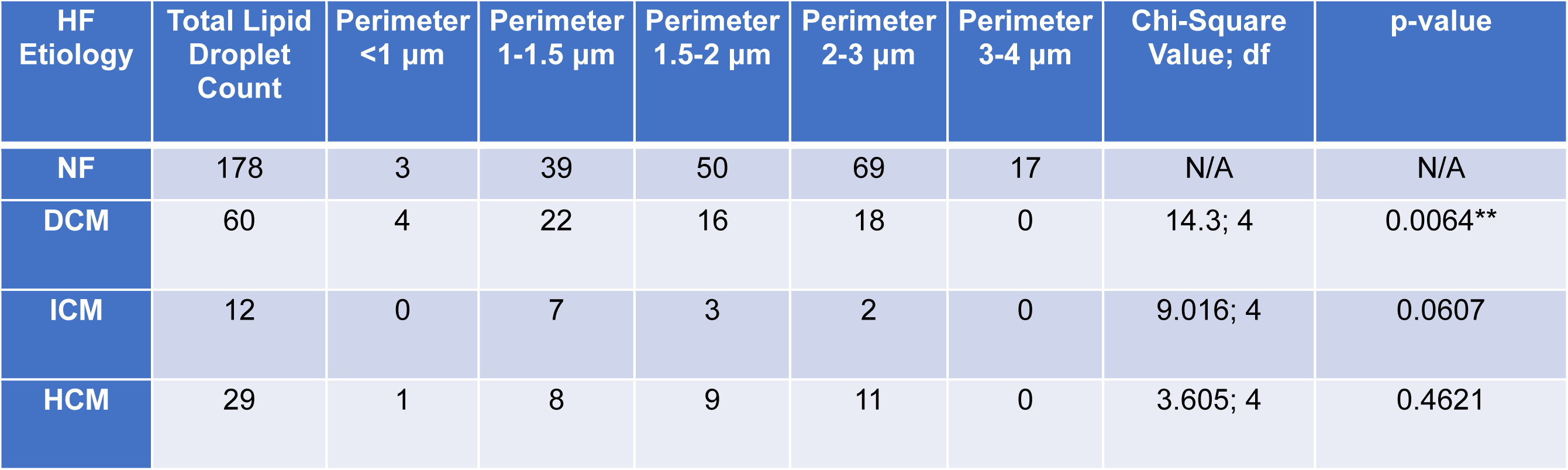
Lipid Droplet (LD) count and LD perimeter size from different heart failure etiologies used in figure 3. Chi-Square value and df comparing non-failing group to corresponding HF etiology. Abbreviations: HF- Heart Failure, NF-Nonfailing, DCM- Dilated Cardiomyopathy, ICM- Ischemic Cardiomyopathy, HCM- Hypertrophic Cardiomyopathy.

### Lipid Droplet-Mitochondria association is diminished in failing cardiomyocytes

Lipid droplets and mitochondria are physically and functionally linked to meet the high energetic demand of cardiomyocytes, where this interaction facilitates the transfer of fatty acids to mitochondria, which then undergo β-oxidation to generate ATP for cardiomyocyte function^36–38^. While there is a growing appreciation for the role of lipid droplets and lipid droplet-organelle interaction, the role of lipid droplet-organelle interaction in cardiomyocyte homeostasis remains understudied ^36–38^. Given the observed changes in LD ultrastructure and the importance of LD-mitochondria association for lipid transfer and energy homeostasis, we aimed to characterize LD-mitochondria interaction/association in human HF by ultrastructure analysis (Figure 4A-B). While no significant differences were observed in mean LD-mitochondria distances within 100 nanometers between groups, we did observe a significant decrease in the minimum LD-mitochondria distance in DCM versus NF cardiomyocytes (Figure 4C-D, S4A-B). Further analysis revealed both the length of OMM perimeter in contact with LDs and the length of LD perimeter in association with mitochondria within 100nm was decreased across all HF etiologies, suggesting disrupted LD-mitochondrial interactions in heart failure (Figure 4E-F, S4C-D). When LD-OMM distances were binned, a downward shift in LD perimeter in association with OMM was observed in cardiomyocytes with DCM (Figure 4G, S4E), suggesting that LDs in proximity to mitochondria may have impaired lipid storage or increased lipolysis. While there was a significant decrease in the length of LD-mitochondria associations, there were no significant differences in the number of LD contacts between failing and nonfailing cardiomyocytes (Figure 4H, S4F). Overall, this suggests less functional interaction between mitochondria and LDs in all etiologies of HF. To further investigate any potential mechanisms that may contribute to changes in LD-mitochondria association, we performed qPCR analysis on the human LV tissues for known or suspected LD-mitochondrial tethers. No significant changes in the expression of proposed LD-mitochondria tethers *PLIN3*, *MIGA2*, *FACL2*, or *SNAP23* were observed between groups (Figure 4I-L). Overall, our data suggests that there is impaired lipid-mitochondria association in failing cardiomyocytes and that canonical LD tethers are likely not implicated.

**Figure 4.**
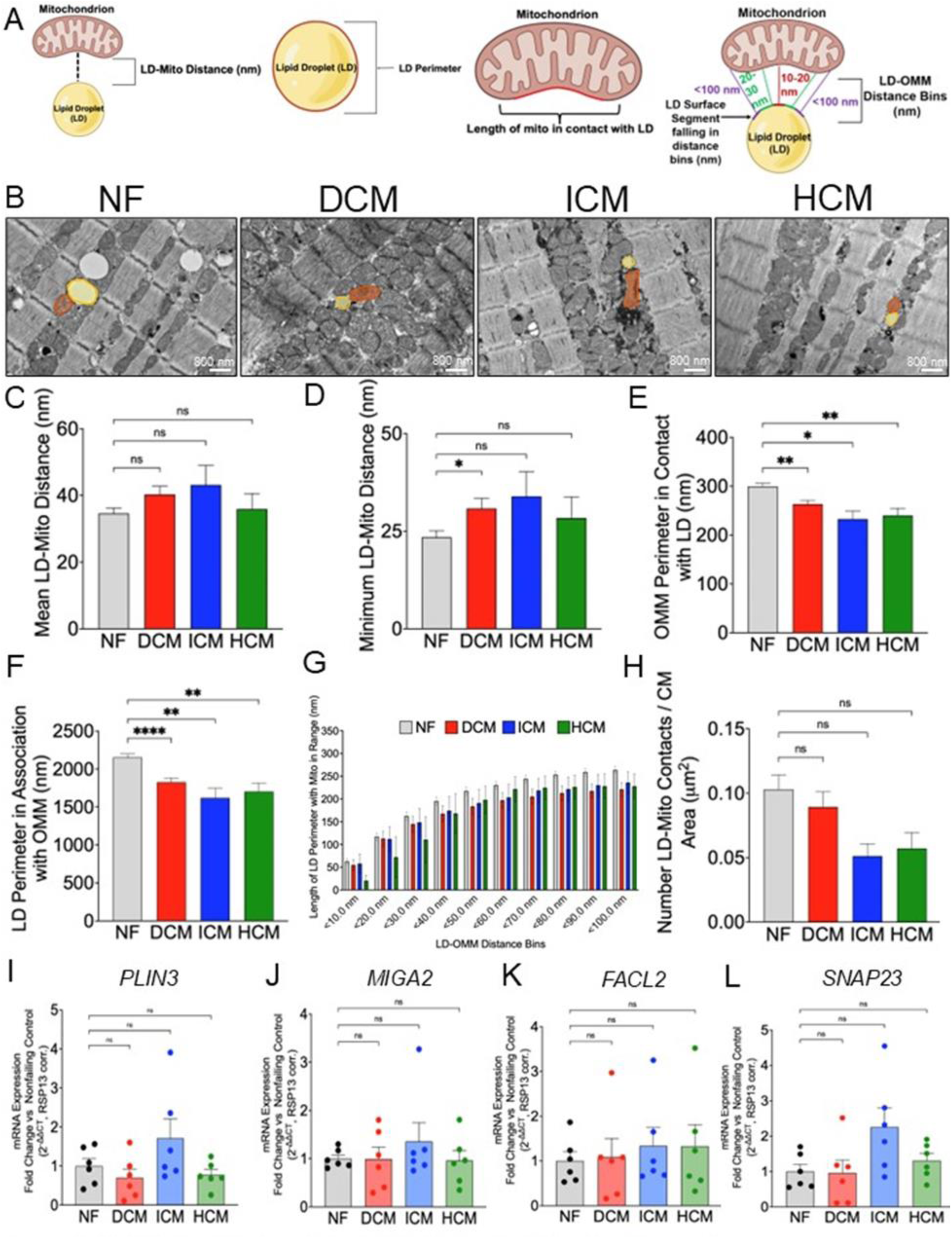
LD-Mitochondria Association is diminished in failing cardiomyocytes. (A). Schematic of LD-mitochondria apposition analysis. (B) Representative EM Images. Cardiac tissue from NF, DCM, HCM, ICM samples were sectioned and imaged by TEM. TEM images were analyzed and quantified using ImageJ Fiji (Scale bars: 800nm). Ultrastructure measurements include (C) mean LD-mitochondria distance, (D) minimum LD-Mito distance, (E) LD perimeter within 100nm of OMM, (F) OMM perimeter in contact with LD, (G) Binning of different LD perimeter sizes within 100nm of OMM in different HF etiologies, (H) Total LD-Mito contacts within image area, (I) mRNA expression of proposed LD-Mito tethers including *PLIN3*, (J) *MIGA2*, (K) *FACL2*, (L) and *SNAP23.* Number of patients=8 NF, 9 DCM, 3 ICM, 5 HCM Number of contacts=60 NF, 33 DCM, 8 ICM, 12 HCM. Number of patients for qPCR = 6 NF, 6 DCM, 6 ICM, 6HCM. Individual data points presented in supplemental. Data is presented as mean ± SEM. ***p<0.001, **p<0.01, *p<0.05; One-way ANOVA with Dunnett’s Post Hoc Test unless stated otherwise. 4G Two-Way ANOVA with Dunnett’s Post Hoc Test. Abbreviations: EM: Electron Microscopy, HF- Heart Failure, NF-Nonfailing, DCM- Dilated Cardiomyopathy, ICM- Ischemic Cardiomyopathy, HCM- Hypertrophic Cardiomyopathy, TEM-Transmission Electron Microscopy, OMM- Outer Mitochondria Membrane, Mito- Mitochondria, *PLIN3- Perilipin 3, MIGA2- Mitoguardin-2, FACL2- Acyl-CoA Synthetase Long Chain Family Member 1, SNAP23- Synaptosome Associated Protein 23*.

## DISCUSSION

The organization and morphology of mitochondria play a critical role in cardiomyocyte function. In cardiomyocytes, mitochondria tend to be elongated and are organized between myofibrils. This organization maximizes the efficiency of calcium signaling necessary for excitation-contraction coupling and ATP generation^4–17,22^. Changes in the shape, size and organization of mitochondria have been reported to impact various mitochondrial functions including biogenesis, calcium signaling and apoptosis^78–84^. While changes in cardiac ultrastructure in HF have been previously reported, how organelle apposition is perturbed in human heart failure remains poorly understood^4–17,24–48^. In this study we characterized organelle apposition in human heart failure to try and define clinical significance.

Here, morphometric analysis of mitochondria ultrastructure revealed failing cardiomyocytes had smaller and more circular mitochondria compared to nonfailing controls. These smaller and more fragmented mitochondria are a key indicator of abnormal mitochondrial dynamics and mitochondrial dysfunction and is suggestive of increased fission^12–18^. Mitochondria are double membrane organelles; the folds of the inner membrane are referred to as cristae which play a critical role in ATP generation and mitochondrial signaling^6–18,79–85^. Mitochondria size is known to impact ATP production, suggesting the smaller, circular mitochondria observed in failing cardiomyocytes may lead to reduced membrane surface area and disrupted cristae organization necessary for appropriate bioenergetics. Disruptions in cristae organization is associated with a reduction in the number of ATP synthase complexes and electron transport chain components, which could cause a decrease in ATP production^6–18,79–85^. Taken together, the observed changes in mitochondria morphology in failing cardiomyocytes have several implications including impaired calcium homeostasis, diminished bioenergetics, and disrupted mitochondrial dynamics, all processes critical for cardiomyocyte function and contractility.

To ensure cardiac health, mitochondria orchestrate a complex signaling network, integrating energy signaling to meet the energetic demands of the cell. Additional critical signaling functions of the mitochondria in the cardiomyocyte include calcium homeostasis, reactive oxygen species (ROS) production, organelle communication and mitochondrial dynamics^7–22^. While the generation of ATP is a critical function of mitochondria to provide a stable source of energy to the cardiomyocyte, ATP also regulates the activation of AMP-activated protein kinase (AMPK)^7–16^. AMPK signaling in the heart is a critical mediator of cardiac metabolism, but also regulator of protein synthesis and various stress responses linked to heart failure^4–8,11–22^. Mitochondria are also essential for buffering calcium released from the SR, allowing for efficient ECC coupling and linking bioenergetics to cardiac contractions^4–8,11–22^. These calcium signals regulate mitochondrial dehydrogenases in the TCA cycle, ensuring metabolic activity matches the cardiac demands^4–8,11–22,29–32^. Mitochondria are also major source of ROS which at physiological levels promote cell survival and regulates hypertrophic signaling^14–19^. However, excessive ROS production results in oxidative damaging furthering heart failure procession^14–19^. Impairments in the number or function of mitochondria can perturb both energy and ROS production, highlighting the importance of appropriate mitochondrial fission and fusion dynamics to maintain cardiac homeostasis^4–8,11–22,29–32^.

Changes in mitochondria morphology are regulated by the processes of fission and fusion^12–18^. Mitochondrial fusion helps to maintain mitochondrial homeostasis and lessen the effects of damage by mixing the contents of individual mitochondrial into a single compartment. Mitochondrial fission splits mitochondria into two separate organelles aiding in the proper distribution of mitochondria during cell division and maintaining quality control by removing damaged mitochondria via autophagy or mitophagy^12–18,78–84^. Imbalances in these processes result in either fragmented or large fused mitochondria, respectively, which can result in disrupted mitochondrial function and perturbed cardiac function^12–18,78–84^. Our study revealed an increased number of smaller mitochondria in failing cardiomyocytes, suggesting an imbalance in mitochondria dynamics favoring fission^12–16^. Several proteins are responsible for maintenance of mitochondrial morphology. Mitofusin 2 (MFN2) and optic atrophy factor 1 (OPA1) mediate mitochondrial fusion, while dynamin-related protein 1 (DRP1), and human fission factor-1 (FIS1) mediate fission^12–18,75–85^. While altered expression of various tethers in different forms of HF were identified, no consistent changes were shared between all studied forms of HF. These tethering proteins have been shown to have complex post-translational modifications, which may explain why SR-mitochondrial proximity was consistently disrupted in all forms in HF despite varying changes in mRNA expression. Recent studies have highlighted mitochondrial dynamics as a promising therapeutic target. A study utilizing a cell-permeant peptide to modulate mitochondrial fusion by targeting *Mfn2* conformation was shown to correct mitochondria abnormalities in cultured fibroblasts and neurons with Charcot-Marie-Tooth disease type 2A (CMTA2) defects^82^. It has also been shown that when Elamipretide, a small peptide that binds to cardiolipin on mitochondria, was employed it demonstrated a reduction in mitochondrial fragmentation in the fibroblasts of patients with dilated cardiomyopathy with ataxia syndrome^87^. Furthermore, in the context of cardiac ischemia-reperfusion, the DRP1 inhibitor (mdivi-1) was shown to inhibit fission and promote cardioprotection^16^. Gaining a deeper understanding of the molecular machinery modulating mitochondrial morphology can provide novel targets for HF treatment.

Organelle apposition is instrumental for communication between organelles and is a critical regulator of intracellular calcium (_i_Ca^2+^) exchange, lipid transfer, mitochondrial dynamics and autophagy^23–38^. Emerging evidence has identified disrupted organelle communication in cardiac pathology, including alterations in SR-mitochondrial tethering which impacts cardiac metabolism and calcium handling, contributing to cardiovascular pathologies including heart failure^24–33,40–48^. Gain- and loss-of-function studies in murine models of *Mfn2* and protein tyrosine phosphatase interacting protein 51 (*Ptptip51*) demonstrated increases and decreases in SR-mitochondria tethering^65^. *Mnf2* cardiac knockout mice demonstrated decreased mitochondrial contact length with the junction SR and a cardiac phenotype of disrupted mitochondrial bioenergetics, as well as mild hypertrophy and decreased cardiac function^40,63–64^. Knockdown of *Ptpip51* improved heart function following ischemia-reperfusion injury^65^. A recent study by Csordas *et al.* demonstrated that the use of a synthetic linker, which maintains close proximity between the junctional SR and mitochondria, reduced cardiomyocyte death and attenuated calcium-overload in an *ex vivo* ischemia-reperfusion injury model^41^. Overall, the synthetic linker improved cardiac contractility and protected female mice from early decompensation induced by adrenergic stress, further highlighting the significance of SR-mitochondria tethering in cardiac function^41^. Given the importance of SR-mitochondria tethering in cardiac physiology we evaluated if SR-mitochondrial apposition was changed in human HF. We found that cardiomyocytes from patients with DCM, HCM and ICM exhibited a significant increase in SR-mitochondrial distance and decreased functional interaction between these organelles. The decrease in physical proximity may contribute to reduced communication between these organelles, impacting functions critical for cardiomyocyte homeostasis, such as calcium transfer and lipid homeostasis.

Numerous proteins have been implicated in the modulation of the SR-mito tethering complex^23–33,42–69^. Analysis of mRNA expression of proposed tethers in human left ventricular samples revealed significant changes in expression in a number of these components during heart failure, with HCM having the most significant changes in tether expression. This includes the recently discovered Mitofusin splice variant, ERMIT2, which is shown to be enriched at the ER-mitochondria interface and tether the ER to mitochondria^85^. Collectively, this data suggests the expression of tethering components may be perturbed in HF and may contribute to disrupted SR-mitochondria crosstalk seen in failing cardiomyocytes. This provides clinically relevant evidence to further study the implications of SR-mitochondria contacts in human heart failure.

Lipid droplets (LDs) are dynamic organelles that fluctuate in size to meet the energetic demand of the cell^34–36,86^. Their functions include the storage of fatty acids, maintenance of lipid homeostasis and membrane maintenance. Despite their significance in critical cellular processes, LD size, morphology, and localization in relation to mitochondria have remained poorly characterized across biology, including HF^34–36,86^. Interestingly, our analysis revealed a significant reduction in the number and size of LDs present in failing cardiomyocytes. This could be explained by potentially more lipolysis that disrupts lipid storage and metabolomic homeostasis. While lipolysis may initially be beneficial by preventing LD accumulation, prolonged lipolysis can contribute towards perturbed energetics and promote HF^34–36,86^. Further research is required to understand the mechanisms modulating LD size in HF and the causal implications.

Cardiomyocytes require high levels of energy compared to other tissue types due to their role in maintain cardiac output. To meet this energetic demand, mitochondria utilize fatty acids as their primary source to generate ATP^7–16,34–38^. This process requires close association and proximity between LDs and mitochondria to enable the efficient transfer of lipids to mitochondria for β-oxidation for ATP generation^34–38,70–77^. Our analysis identified reduced LD-mitochondria association in failing cardiomyocytes compared to non-failing cardiomyocytes. This decrease in functional interplay between LDs and mitochondria may potentially disrupt fatty acid transfer between the organelles and impair energy production. Several proteins have been implicated in maintaining LD-mitochondria interactions including PLIN5 and mitochondria-associated protein MIGA2^16,70–77^. Perilipins are located on the surface of lipid droplets and proteins within in the perilipin family have been shown to play a role in the regulation of LDs. While PLIN5 has linked lipid storage to mitochondria function, Perilipin 3 (PLIN3) has been implicated in lipid droplet dynamics^16,70–71^. Another proposed LD-mitochondria tether, MIGA2 has been shown to mediate the ER-mitochondria contact sites and facilitate lipid transfer^72–73^. While we did not observe any significant change in proposed LD-mitochondria tethers at the mRNA level, it remains possible that post-translational modifications and protein stability may still be altered in HF that could explain the differences we identified during ultrastructure analysis. Overall, our findings emphasize the importance of LD dynamics and LD-mitochondria juxtaposition in human HF pathophysiology, highlighting the need for causal research to investigate how LD-mitochondria communication impacts cardiac energy metabolism.

Although this study provides novel insight into organelle apposition it is not without limitations. The use of human cardiac tissues from end-stage failing human hearts is limited to availability and may result in variability due to disease etiology and treatment. This study utilized TEM to provide basic morphometric analysis of organelles and characterization organelle apposition. Future studies employing FIB-SEM will allow for investigation of dynamic organelle interactions such as those between the SR-mitochondria and LD-mitochondria and allow for quantification of interaction volume. In addition, understanding how protein tethers impact organelle apposition and how SR-mitochondria contacts impact mitochondria morphology requires investigation. Further research is needed to uncover the mechanisms that govern the physical and metabolic interactions between cardiac lipid droplets and mitochondria, as well as their physiological and pathological significance. While our study looked extensively at mRNA expression patterns of various tethers, future work will focus on evaluating protein expression and post-expression modifications that may contribute towards perturbed tethering. Finally, causal examination of specific tethers is needed to define how these processes impact overall HF progression.

In summary, through comprehensive TEM analysis, we show that failing human cardiomyocytes exhibit both altered mitochondrial and lipid dynamics compared to nonfailing cardiomyocytes. Increased mean SR-mitochondrial distance and decreased SR-mitochondrial association were also observed in failing cardiomyocytes. Our findings suggest a role in organelle apposition in the progression of heart failure pathophysiology, providing rationale for future studies aimed at uncovering the mechanisms organelle apposition. Collectively, these studies underscore the importance of uncovering mechanisms contributing to the structural defects and governing organelle tethering. Understanding the disruption of organelle apposition offers novel insights into HF pathogenesis and could lead to the identification of potential therapeutic targets

## ACKNOWLEDGMENTS

We would like to thank Ken Bedi and Dr. Ken Margulies (UPenn). We also thank S. Modla and J. Ross for her expertise in TEM sample processing (Delaware Biotechnology Institute).The authors thank Dr. Gyorgy Csordas and Dr. Zuzana Nichtova at MitoCare at Thomas Jefferson University for their expertise in TEM imaging and analysis.

## AUTHOR CONTRIBUTIONS

Conception and design of research: N.R.L., and J.W.E.

Performed experiments: N.R.L., K.C.B., and B.L.P.

Analyzed data: N.R.L., T.L.S and J.W.E.

Interpreted results of experiments: N.R.L., T.L.S and J.W.E.

Prepared figures: N.R.L.

Drafted manuscript: N.R.L.

Edited and revised manuscript: N.R.L., T.L.S and J.W.E.

## SOURCES OF FUNDING

National Institutes of Health grant R01NS121379 (JWE)

National Institutes of Health grant P01HL134608 (JWE)

National Institutes of Health grant P01HL147841 (JWE)

National Institutes of Health grant R01HL142271 (JWE)

American Heart Association award 20EIA35320226 (JWE)

National Institutes of Health grant T32HL091804 (JWE, TLS)

## DISCLOSURE

Authors have no disclosures.

## Non-standard Abbreviations and Acronyms

BAP31: B Cell Receptor Associated Protein 31
Ca^2+^: Calcium
CM: Cardiomyocyte
CMTA2: Charcot-Marie-Tooth disease type 2A
DCM: Dilated cardiomyopathy
DJ1: Parkinsonism Associated Deglycase
DRP1: Dynamin-related protein 1
ECC: Excitation Contraction Coupling
ERMIN2: ER mitofusin 2
ERMIT2: ER mitofusin 2 tether
FACL2: Acyl-CoA Synthetase Long Chain Family Member 1
FACL4: Acyl-CoA Synthetase Long Chain Family Member 4
FIS1: Fission, Mitochondrial 1
FUNDC1: FUN14 domain-containing protein 1
GRP75: Glucose regulated protein 75
HF: Heart failure
HCM: Hypertrophic cardiomyopathy
ICM: Ischemic cardiomyopathy
IP3R1: Inositol 1,4,5-Trisphosphate Receptor Type 1
IP3R2: Inositol 1,4,5-Trisphosphate Receptor Type 2
IP3R3: Inositol 1,4,5-Trisphosphate Receptor Type 3
jSR: Junctional Sarcoplasmic Reticulum
LD: Lipid Droplet
LV: Left Ventricular
MFN2: Mitofusin 2
MIGA2: Mitoguardin-2
NF: Nonfailing
OMM: Outer Mitochondria Membrane
OPA1: OPA1 Mitochondrial Dynamin Like GTPase
ORMDL3: ORMDL Sphingolipid Biosynthesis Regulator 3
PACS2: Phosphofurin Acidic Cluster Sorting Protein 2
PDZD8: PDZ Domain Containing 8
PLIN3: Perilipin 3
PLIN5: Perilipin-5
PTPIP51: Protein tyrosine phosphatase-interacting protein 51
ROS: Reactive Oxygen Species
SIGMAR-1: Sigma Non-Opioid Intracellular Receptor 1
SNAP23: Synaptosome Associated Protein 23
SR: Sarcoplasmic Reticulum
TCA: Tricarboxylic Acid Cycle
TEM: Transmission electron microscopy
TPMS: Trichoplein Keratin Filament Binding
T-Tubule: Transverse tubule
VAPB: Vesicle-associated membrane protein-associated protein B
VDAC1: Voltage Dependent Anion Channel 1
VDAC2: Voltage Dependent Anion Channel 2

## Supplemental Figure Legends

**Supplemental Figure 1.**
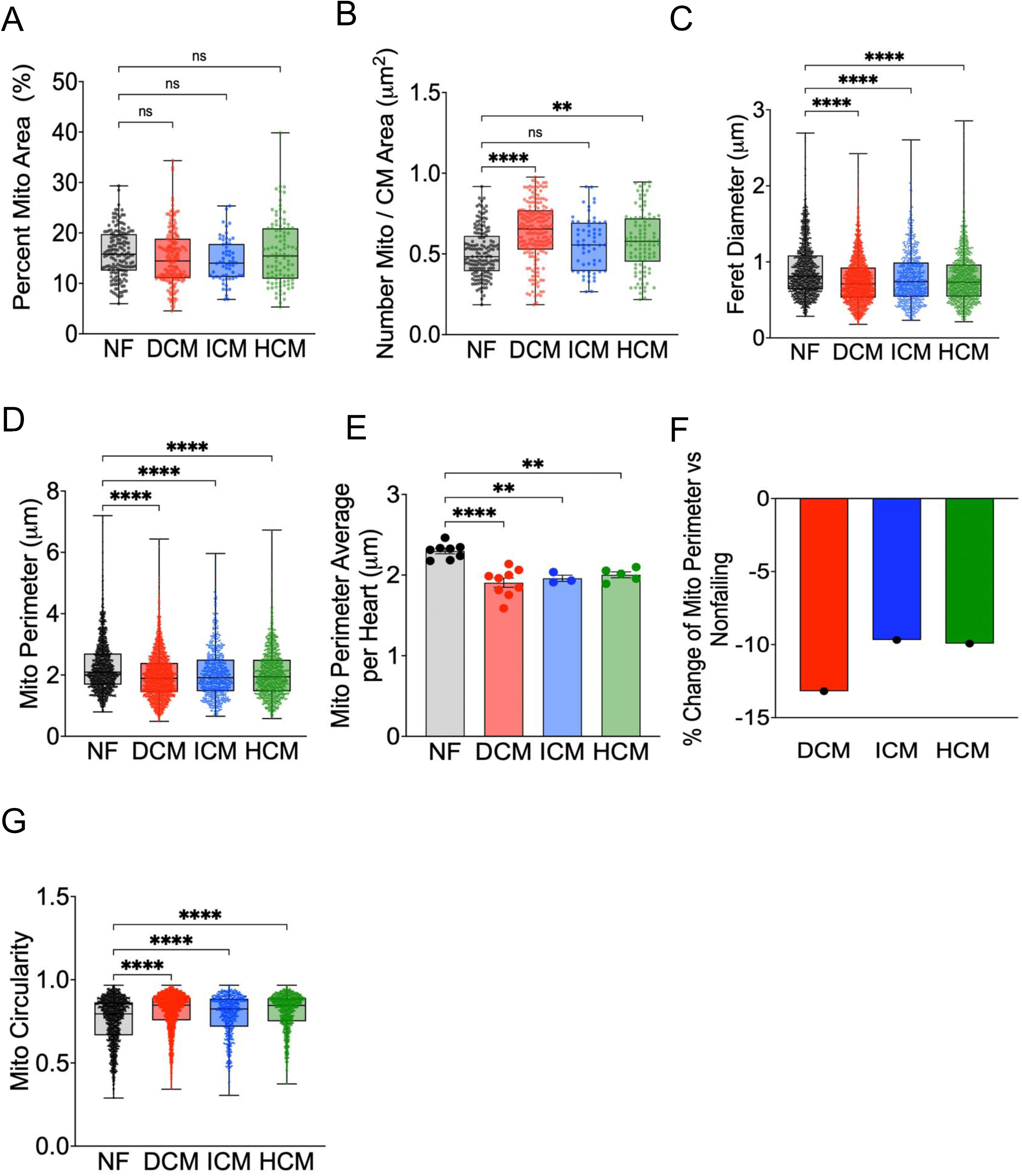
Mitochondrial dynamics is altered in failing human cardiomyocytes. Individual values for ultrastructure measurements include (A) percent mitochondria area, (B) number of mitochondria/cardiomyocyte (CM) area, (C) % change of mitochondria perimeter (D) mitochondria perimeter average per heart, (E) mitochondrial perimeter in HF etiologies, (F) mitochondria feret diameter and (G) mitochondrial circularity. Number of patients for EM Data=8 NF, 9 DCM, 3 ICM, 5 HCM. Number of mito= 2760 NF, 4546 DCM, 1368 ICM, 2350 HCM. Data is presented as mean ± SEM. ****p<0.0001, **p<0.01, *p<0.05; One-way ANOVA with Dunnett’s Post Hoc Test.

**Supplemental Figure 2.**
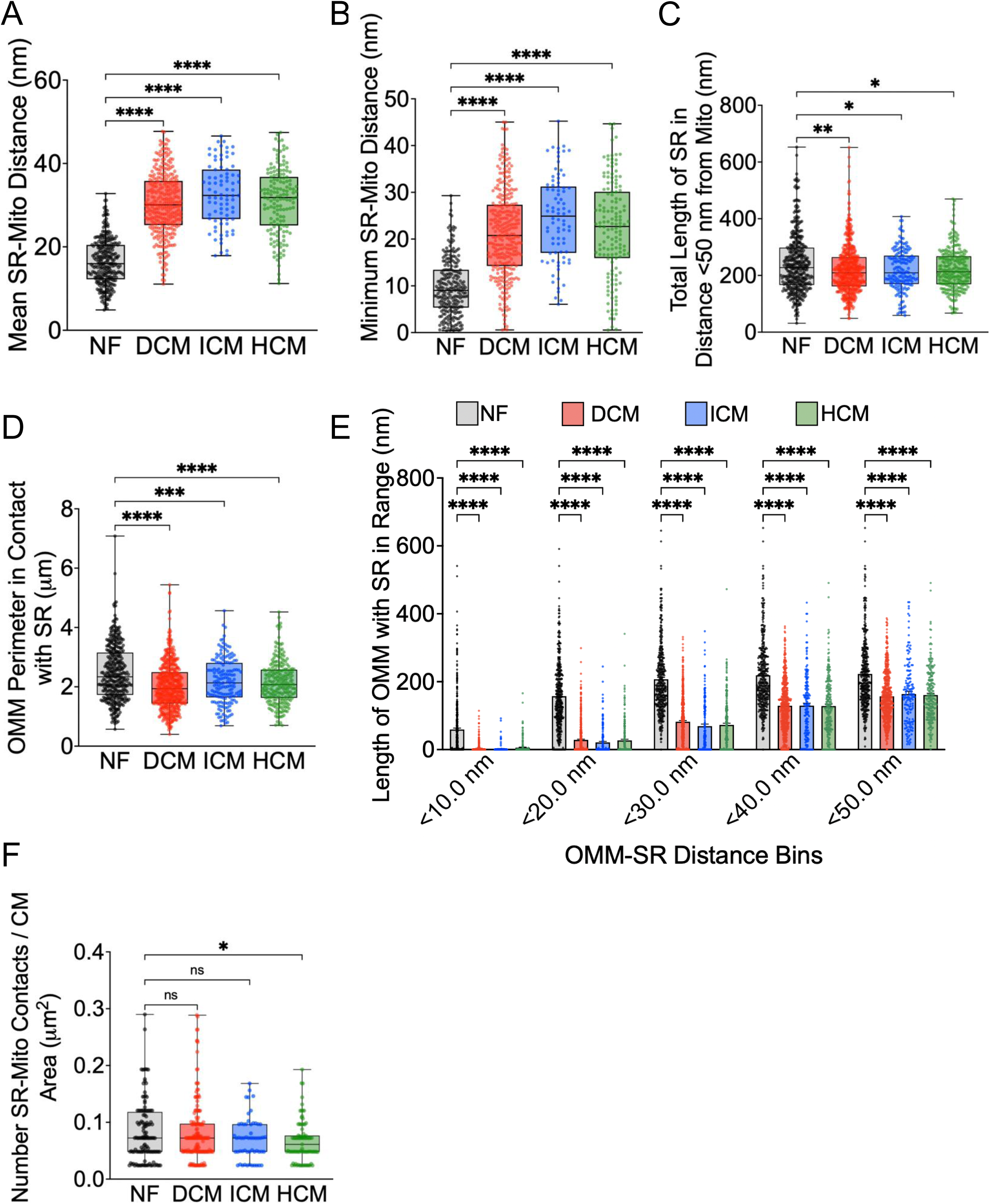
Failing cardiomyocytes have altered sarcoplasmic reticulum-mitochondria apposition. Individual values for ultrastructure measurements include (A) mean SR-mitochondria distance, (B) minimum SR-mitochondria distance, (C) total length of SR less than 50 nm from mitochondria, (D) mitochondrial perimeter in association with SR, (E) binning of the length of OMM less than 50 nm from SR, (F) SR Length/Mito Perimeter, (G) and total number of SR-mitochondrial contracts per image area. Number of patients=8 NF, 9 DCM, 3 ICM, 5 HCM Number of contacts=405 NF, 526 DCM, 163 ICM, 237 HCM. Data is presented as mean ± SEM. ****p<0.0001, **p<0.01, *p<0.05; One-way ANOVA with Dunnett’s Post Hoc Test unless stated otherwise. 2E Two-Way ANOVA with Dunnett’s Post Hoc Test

**Supplemental Figure 3.**
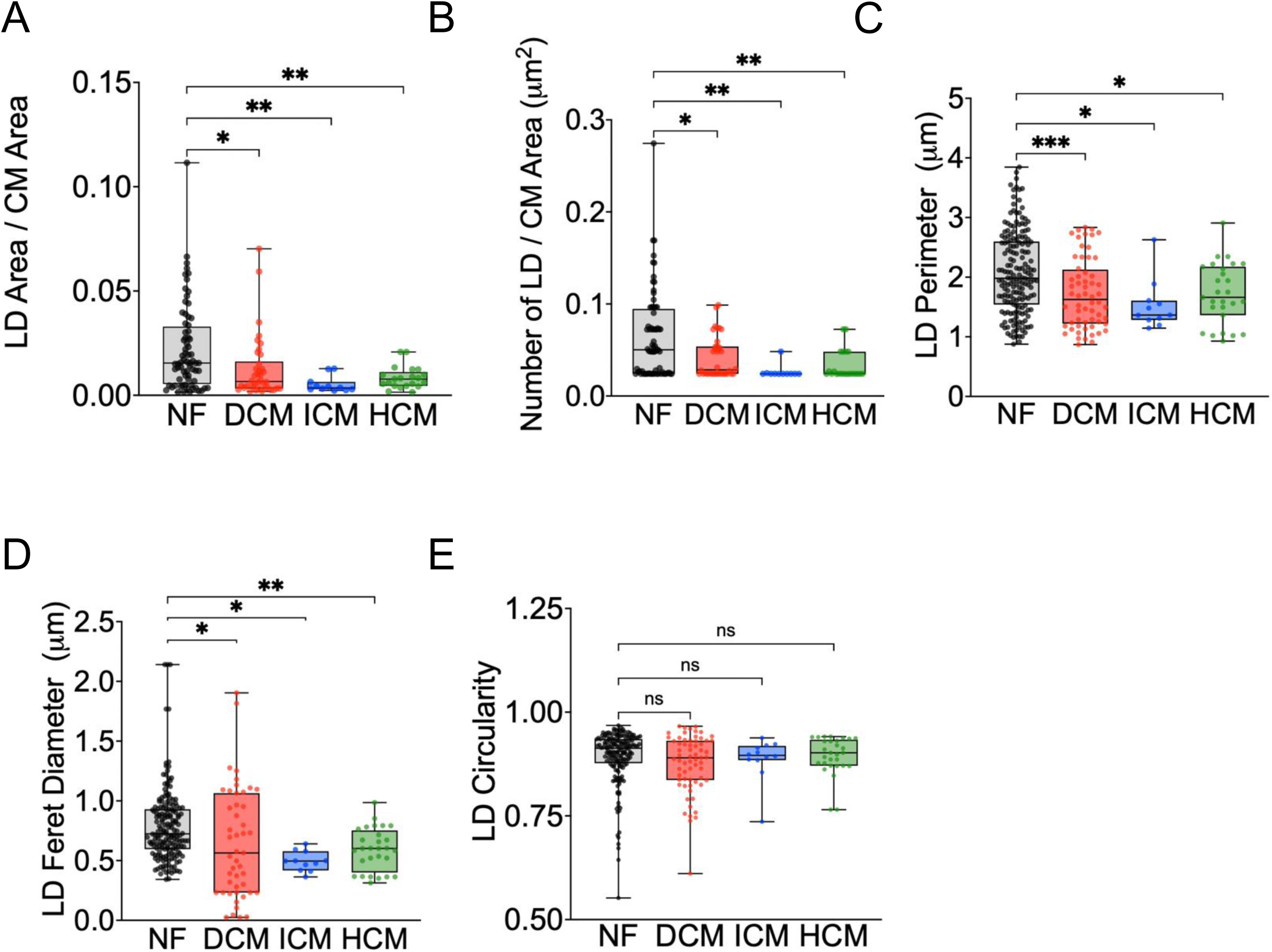
LD ultrastructure is altered in failing cardiomyocytes. Individual values for ultrastructure measurements include(A) LD area/total cardiomyocyte area, (B) number of LD/CM area, (C) LD perimeter, (D) LD Feret diameter, (E)and LD Circularity. Number of patients=8 NF, 9 DCM, 3 ICM, 5 HCM Number of LD= 178 NF, 60 DCM, 12 ICM, 29 HCM. Data is presented as mean ± SEM. **p<0.01, *p<0.05; One-way ANOVA with Dunnett’s Post Hoc Test.

**Supplemental Figure 4.**
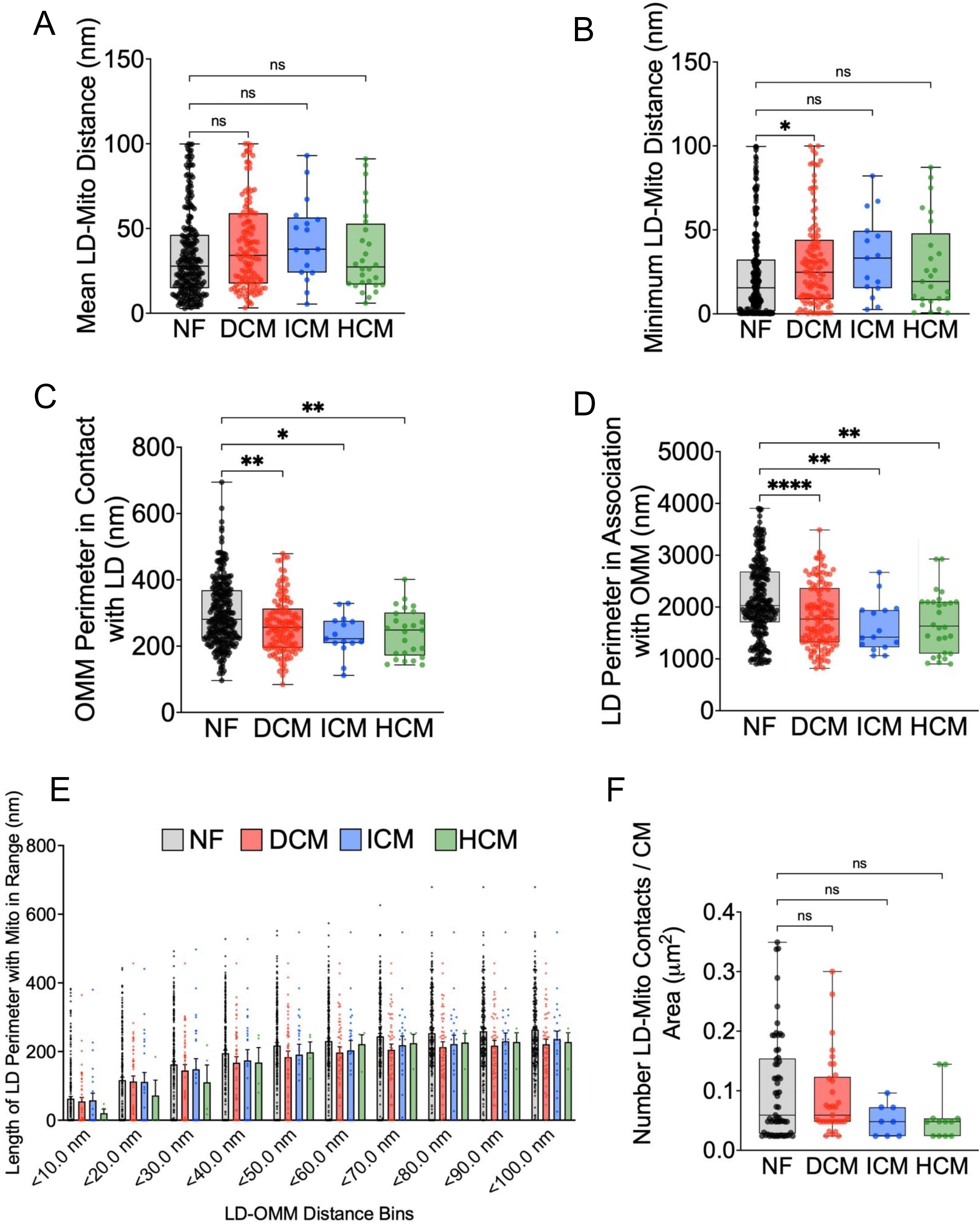
LD-Mitochondria Association is diminished in failing cardiomyocytes. Individual values for ultrastructure measurements include (A) mean LD-mitochondria distance, (B) minimum LD-Mito distance, (C) OMM perimeter in contact with LD (D) LD perimeter within 100 nm of OMM,(E) Total LD-Mito contacts within image area and (F) Binning of different LD perimeter sizes within 100nm of OMM in different HF etiologies, Number of patients=8 NF, 9 DCM, 3 ICM, 5 HCM Number of contacts=60 NF, 33 DCM, 8 ICM, 12 HCM. Number of patients for qPCR = 6 NF, 6 DCM, 6 ICM, 6HCM. Data is presented as mean ± SEM. ***p<0.001, **p<0.01, *p<0.05; One-way ANOVA with Dunnett’s Post Hoc Test unless stated otherwise. 4F Two-Way ANOVA with Dunnett’s Post Hoc Test

## Notes

### Competing Interest Statement

The authors have declared no competing interest.

